# A Shared Pathogenic Mechanism for Valproic Acid and SHROOM3 Knockout in a Brain Organoid Model of Neural Tube Defects

**DOI:** 10.1101/2023.04.11.536245

**Authors:** Taylor N. Takla, Jinghui Luo, Roksolana Sudyk, Joy Huang, J. Clayton Walker, Neeta L. Vora, Jonathan Z. Sexton, Jack M. Parent, Andrew M. Tidball

## Abstract

Neural tube defects (NTDs) including anencephaly and spina bifida are common major malformations of fetal development resulting from incomplete closure of the neural tube. These conditions lead to either universal death (anencephaly) or life-long severe complications (spina bifida). Despite hundreds of genetic mouse models having neural tube defect phenotypes, the genetics of human NTDs are poorly understood. Furthermore, pharmaceuticals such as antiseizure medications have been found clinically to increase the risk of NTDs when administered during pregnancy. Therefore, a model that recapitulates human neurodevelopment would be of immense benefit to understand the genetics underlying NTDs and identify teratogenic mechanisms. Using our self-organizing single rosette spheroid (SOSRS) brain organoid system, we have developed a high-throughput image analysis pipeline for evaluating SOSRS structure for NTD-like phenotypes. Similar to small molecule inhibition of apical constriction, the antiseizure medication valproic acid (VPA), a known cause of NTDs, increases the apical lumen size and apical cell surface area in a dose-responsive manner. This expansion was mimicked by GSK3β and HDAC inhibitors; however, RNA sequencing suggests VPA does not inhibit GSK3β at these concentrations. Knockout of SHROOM3, a well-known NTD-related gene, also caused expansion of the lumen as well as reduced f-actin polarization. The increased lumen sizes were caused by reduced cell apical constriction suggesting that impingement of this process is a shared mechanism for VPA treatment and SHROOM3-KO, two well-known causes of NTDs. Our system allows the rapid identification of NTD-like phenotypes for both compounds and genetic variants and should prove useful for understanding specific NTD mechanisms and predicting drug teratogenicity.

## INTRODUCTION

Open neural tube defects (NTDs) are common congenital malformations (∼1/1000 live births in the US) including anencephaly and spina bifida^1^. These major malformations of neurodevelopment lead to spontaneous abortion or neonatal death in the case of anencephaly, and paraplegia or severe physical or cognitive disabilities for spina bifida. These malformations arise from genetic defects or environmental exposures, including toxins, pharmaceuticals, and nutrient deprivation. The need for robust teratogen screening of novel therapeutics was highlighted by the thalidomide tragedy of the 1950s. This compound was marketed as a treatment for morning sickness during pregnancy. Although rodent testing did not indicate teratogenic risk of thalidomide use, this drug unexpectedly led to a dramatic number of congenital malformations, including NTDs, in human fetuses^2^. Despite this tragedy, rodent models remain the standard method for teratogenic screening, even though the problem of species-specific differences persists. Overall, rodent models of drug toxicity have shown a low rate of concordance with humans leading to the termination of many clinical trials due to human-specific toxicities^3^, and neurological toxicities are the most common cause of clinical trial termination (22%). In addition to the species-specific problems with current teratogenicity screening, these models are costly, labor intensive, and result in a moderate rate of false positives and false negatives^4^. For these reasons, President Biden signed the “FDA Modernization Act 2.0” on December 29, 2022 which removes the mandate for animal testing for new drugs and specifically states that alternative testing methods should include “cell-based assays, organ chips, and microphysiological systems”^5^.

The first pharmaceutical known to produce congenital malformations in humans was the chemotherapy drug, aminopterin^6^ that blocks the folic acid pathway. Folic acid deficiency is the most prominent identified risk factor of congenital malformations^7^. Other commonly used pharmaceuticals associated with NTDs are valproic acid (VPA), primarily an antiseizure medication, lithium, a mood stabilizer used for bipolar disorder and major depressive disorder, and possibly HIV antiretroviral therapies^8–11^. Women taking VPA have a 10-fold increased risk of NTD pregnancies^8, 12^. Since its therapeutic discovery, VPA has been found to act on several molecular targets, including as an inhibitor of embryonic folate metabolism^13^, histone deacetylases (HDACs)^14^ and glycogen synthase kinase-3 (GSK3)^15^, and by increasing oxidative stress^16^. However, the precise teratogenic mechanism(s) of VPA remains unknown.

Open NTDs are also known to have a large genetic risk theoretically due to rare *de novo* or inherited deleterious genetic variants. This genetic risk is shown by a higher concordance of NTD for monozygotic twins (7.7%) than dizygotic twins (4.0%)^17, 18^. Additionally, after a single NTD pregnancy, the risk of recurrence goes from ∼1/1000 to 1/20. Despite this accumulating evidence for a large genetic risk component and >350 gene genes known to cause NTDs in knockout mice, little is known about the genetic variants that cause NTDs in humans. This is likely due to complex genetics, incomplete penetrance, gene-environment interactions (e.g. folic acid), and lack of information on the effects of specific mutations (e.g. missense)^19^. *SHROOM3* loss-of-function variants have been shown to cause exencephaly in genetic mouse models and have been identified in human open NTD pregnancies^20–24^. *SHROOM3* is an important scaffold protein for apical-basal cell polarity organization^25^. It coordinates apical constriction via polarized recruitment of rho-kinase which leads to phosphorylation of non-muscle myosin and, ultimately, constriction of the apical actomyosin cytoskeletal network. This polarized apical constriction is necessary to initiate the process of neurulation. In addition to rare or *de novo* germline mutations, a recent hypothesis for the sporadic nature and lack of genetic variant identification in many human NTD pregnancies is that somatic mosaic mutations in the neural tube can also lead to an open NTD. One recent study provided evidence for this hypothesis by showing that when only 16% of cells in the mouse embryo neural tube had the NTD gene *VANGL2* knocked out spina bifida resulted^26^. Identification of potential human NTD variants and characterization of these variants in robust human NTD models is needed to provide information for effective genetic counseling and family planning.

Human pluripotent stem cell-based models already provide greater predictive value than animal models for certain types of toxicological screening. For example, proarrhythmic risk is now routinely assessed in iPSC-derived cardiomyocytes ^27^. Not only can these human models avoid false negative results like thalidomide, but may potentially limit false positives, allowing for a larger number of medications to enter clinical trials. With the advent of 3D brain organoid technology, several groups have made attempts to model neuroteratogenicity ^28^. Unfortunately, standard methods result in highly variable organoids with multiple-rosettes^28^. Since neural rosettes are the *in vitro* correlate of the developing neural tube, a multirosette model does not recapitulate normal human brain development adequately for detailed structural analyses. Therefore, most models have used a transcriptomic approach to assess the neuroteratogenicity of compounds such as the STOP-Tox_UKN_ system^29^ or involved complicated machine learning algorithms to identify and measure 2-dimensional structures in the neural rosette formation assay^30^. Attempts have also been made to model genetic NTDs as well^31, 32^. One study compared brain organoids from spina bifida aperta patients with controls finding differences in the size and number of neural rosettes between disease groups^32^. However, this study lacked isogenic controls, and iPSCs were generated from different tissue types. A more recent study modeled recurrent anencephaly due to recessive mutations in *NUAK2* with iPSCs and organoids. While reaching statistical significance, the structural organoid measurements were extremely heterogeneous due to multiple rosettes and variable sectioning plane. Furthermore, the structural results were difficult to correlate with mechanism^33^. Given these issues, a 3D brain organoid model that recapitulates neurodevelopment in humans with higher fidelity would be useful to examine the genetic mechanisms of NTDs.

Our group and others have published techniques for generating single rosette organoids or neural cyst cultures^34–38^. Our simple model of human neurulation, which we call SOSRS (self-organizing single-rosette spheroids), overcomes the lack of disease-relevant structural readout in previous brain organoid models. Herein, we demonstrate clear, *in vitro,* dose-responsive structural phenotypes for acute VPA treatment, characterize the transcriptomic effects of VPA, and identify inhibition of apical constriction via HDAC inhibition as the likely mechanism by which VPA causes NTDs. Furthermore, we demonstrate altered structure, polarization, and reduced apical constriction for *SHROOM3* knockout SOSRS, and we find that these phenotypes are gene-dose responsive in mosaic mixing experiments. These findings provide additional evidence that somatic, mosaic variants could lead to NTD formation.

## MATERIALS AND METHODS

### iPSC lines

We purchased the AICS-0023 cell line from the Coriell Institute Biorepository in the Allen Cell collection. This line contains a monoallelic mEGFP-TJP1 (which encodes for the zona occludens 1 [ZO1] protein) to label the lumen in live SOSRS. The *SHROOM3* knockout line and isogenic control lines were generated utilizing a simultaneous CRISPR indel formation and iPS cellular reprogramming technique we previously published^39^. In short, commercially available foreskin fibroblasts were nucleofected with episomal reprogramming vectors^40^ and the pX330 CRISPR plasmid containing a gRNA targeting the 5^th^ exon of the *SHROOM3* gene near a premature stop codon identified in an anencephalic human fetus^22^. Clonal iPSC lines generated by the reprogramming were PCR amplified over the gRNA cut site and Sanger sequenced. One isogenic control and one compound heterozygous frameshift (−1/−14bp) clone were identified and sequenced by NGS to ensure a 1:1 ratio of the two mutant alleles. The gRNA and PCR primer sequences are listed in Table S1.

### iPSC culture

The iPS cells cultures were maintained on Geltrex-coated (1:200 dilution in DMEM/F12) 6-well TC dishes in mTeSR1 medium. When the colonies reached ∼40% confluency, the cultures were incubated with 1 mL of L7 dissociation solution (Lonza) for 2 minutes. The solution was then replaced with mTeSR1, and colonies were mechanically detached with a mini cell scraper. The solution was then pipetted up and down 3-6 times to break colonies into smaller pieces. The solution was then re-plated at a dilution of 1:8 onto newly Geltrex-coated plates.

### SOSRS differentiation

iPSC lines were passaged using Accutase (Innovative cell) and replated onto Geltrex-coated (1:50 dilution in DMEM/F12) 12-well plates at 7-8 x 10^5^ cells/well in mTeSR1 with 10 µM rho-kinase inhibitor (Y- 27632; Tocris, 1254). The following day, or when the cells reached 80-100% confluency, the medium was then changed to 3N (50:50 DMEM/F12:neurobasal with N2 and B27 supplements) without vitamin A with 2 µM DMH1 (Tocris, 4126), 2 µM XAV939 (Cayman Chemical, 13596), and 10 µM SB431542 (Cayman Chemical, 13031) (2 mL of medium per well). 75% of this medium was changed daily with 1 µM cyclopamine (Cayman Chemical, 11321) added beginning on day 1. On day 4, the monolayer was cut into squares using the StemPro EZ passage tool (Gibco). The squares were incubated for 2 minutes with L7 hPSC passaging solution (Lonza). After aspirating the L7 solution, the squares were sprayed off the bottom with 1 mL of preconditioned culture media with a P1000 micropipette. An additional 1 mL of fresh culture medium with the 4 inhibitors was added. Approximately 100 µL of resuspended monolayer squares were then transferred into the wells of black-walled thin-bottom 96-well plates (Greiner µClear or PerkinElmer PhenoPlate) preincubated at 37 °C for 30 minutes with 35 µL of 100% Geltrex solution in each well.

### SOSRS drug treatment

Immediately after plating in 96-well plates, SOSRS were treated with various drugs for 48 hours. Along with VPA (Cayman Chemical, 13033), SOSRS were treated with different HDAC inhibitors, GSK3β inhibitors, folic acid (FA) inhibition or supplementation, and antioxidants. We tested HDAC inhibitors that target different classes: Trichostatin A (TSA; Cayman Chemical, 89730), Nicotinamide (Cayman Chemical, 11127), CI-994 (Cayman Chemical, 12084). GSK3β inhibitors included CHIR99021 (Cayman Chemical, 13122), Bio (Cayman Chemical, 13123), SB-216763 (Cayman Chemical, 10010246), SB-415286 (Cayman Chemical, 10010247), and lithium chloride. FA (Cayman Chemical, 20515) was supplemented, and the FA pathway was inhibited by Aminopterin (Cayman Chemical, 21802). Antioxidants included Vitamin E (±-α-Tocopherol Acetate; Cayman Chemical, 28399), and Resveratrol (*trans*-resveratrol [RV]; Cayman Chemical, 70675). Appropriate amounts of each treatment were mixed with 3N media without vitamin A plus the 4 inhibitors. 100μL of media mixed with the treatment was added per well. SOSRS were treated for 48 hours before being fixed.

### SOSRS Fixing & Staining

SOSRS were fixed at day 6 in 4% paraformaldehyde in PBS for 1 hour at 4°C. Following the fixation, cells were washed with PBS twice for 5 minutes at room temperature. SOSRS were permeabilized in PBS with 0.1% TritonX-100 for 30 minutes followed by a 1-hour incubation in ICC blocking buffer (PBS with 0.05% Tween-20, 5% normal goat serum and 1% BSA) + 0.1% TritonX-100. Primary antibodies were mixed with this same blocking buffer and incubated with SOSRS overnight at 4°C. See Table S2 for antibody catalog numbers and dilutions. The SOSRS were then washed 3 times with PBS containing 0.1% Tween-20 (PBST). Secondary antibodies as well as phalloidin were added to ICC blocking buffer and incubated with SOSRS overnight at 4°C. Cells were washed 1 time with PBST. The SOSRS were then counterstained by incubation with either bis-benzimide or HCS CellMask Deep Red for 1 hour at room temperature, followed by 3 additional PBST washes. Alternatively, when immunolabeling was not necessary during the drug treatment experiments, SOSRS were counterstained directly following fixation and then washed with PBS that did not contain Tween-20. In this case, the constitutive expression of the ZO1-EGFP fusion protein was used for image analysis.

### Manual SOSRS imaging and analysis

Images were obtained on an EVOS Cell Imaging System (ThermoFisher). 4 images of different SOSRS were obtained per well. We excluded SOSRS that were smaller than 100 microns or greater than 200 microns in diameter. Larger SOSRS tended to be fusions from 2 monolayer fragments while smaller SOSRS were from smaller pieces of the monolayer. Each SOSRS was imaged for ZO1-EGFP along with bis-benzamide, creating 8 images total for each well (2 per each SOSRS x 4 SOSRS per well). ImageJ was used to set maximum and minimum thresholds to optimize the contrast between the lumen (ZO1-EGFP or ZO1 immunostaining) or SOSRS (bis-benzamide for DNA) with the surrounding background before creating a binary mask. After using the “fill holes” function, the area, circularity, roundness, solidity, and aspect ratio for each was obtained.

### Automated imaging and analysis

Automated confocal microscopy was utilized to increase throughput. For confocal microscopy, the PBS solution was removed from each well and a fructose glycerol solution was added. This solution increased SOSRS clarity by correcting refractive index and dehydrating the Geltrex hydrogel, causing the SOSRS to move to the bottom of dish allowing one imaging range in the Z-axis across the plate. For imaging, we used the Yokogawa Cell Voyager 8000 automated spinning disc confocal microscope with a 10x dry (NA=0.40) UPLSAPO10X2 objective lens. For each well of the 96-well plates, 9 fields were imaged with 16 Z-sections with 6μm spacing and a 100μm total depth. Laser autofocus was performed by undersurface detection and offset of 90μm into the cell plane. This resulted in imaging the vast majority of SOSRS in each well. The optical sections were used to generate a maximum intensity projection for each field. These images were imported into CellProfiler with a custom pipeline to identify the margin of each SOSRS (utilizing HCS CellMask Deep Red) and the apical lumen (utilizing ZO1-EGFP, ZO1 immunostaining, or phalloidin-Alexa488). Using these masks, intensity and morphometric/shape features were measured resulting in over 400 features (Table S3). The database file was then imported into a custom Python script to join experimental metadata (i.e., drug, concentration, genotype, cell line). The script then checks for differences between conditions in the number of lumens in each SOSRS and differences in SOSRS areas. In control lines under normal media conditions, multi-lumen SOSRS and max project areas > 31,415 µm^2^ are mostly driven by two monolayer fragments fusing. Analysis was restricted to single lumen SOSRS with areas between 7,854-31,415 µm^2^. This corresponds to the 100-200 µm diameter criterion used for manual imaging.

### Random forest predictive modeling

To identify the most informative features for discriminating between conditions (VPA concentration or SHROOM3 genotype), we first scaled and centered each metric. We then utilized the bootstrap random forest model and calculated variable importance metrics with standard 5-fold cross validation. This modeling was performed on JMP Pro 16.

### RNA sequencing experiment

Day 6 brain organoids generated from a male iPSC line (AICS-0023) were treated with either a control vehicle or one of the following 3 compounds: 1) the non-teratogenic compound trichostatin-A; 2) the teratogenic anti-seizure medication VPA at a concentration of either 200 uM or 400 uM; or 3) CHIR99021 at a concentration of 10uM. All treatments were performed in 4 replicates, except for CHIR99021 which was performed in triplicate. Total RNA was isolated using the RNeasy PLUS Universal Kit. SOSRS were solubilized along with the Geltrex hydrogel using Qiazol, and the kit was used according to the manufacturer’s recommendations. Libraries were constructed using NEB polyA RNA ultra II and subsequently subjected to 150 cycles of sequencing on NovaSeq-6000 (Illumina). Adapters were trimmed using Cutadapt (v2.3). FastQC (v0.11.8) was used to ensure the quality of data. Reads were mapped to the reference genome GRCh38 (ENSEMBL)] using STAR (v2.6.1b) and assigned count estimates to genes with RSEM (v1.3.1). Alignment options followed ENCODE standards for RNA-seq [4]. FastQC was used in an additional post-alignment step to ensure that only high-quality data were used for expression quantitation and differential expression. Data were pre-filtered to remove genes with 0 counts in all samples. Differential gene expression analysis was performed using DESeq2, using a negative binomial generalized linear model (thresholds: linear fold change >1.5 or <-1.5, Benjamini-Hochberg FDR (Padj) <0.05). Genes were annotated with NCBI Entrez GeneIDs and text descriptions. Plots were generated using variations of DESeq2 plotting functions and other packages with R (version 3.3.3). Data for all expressed genes comparing between treatment and vehicle controls is given in Tables S4-6.

### Quantifying apical surface areas

SOSRS counterstained and/or immunostained as described above were imaged with an oil 100x objective on a Stellaris 5 inverted confocal microscope. Z-series were set at immediately above and below each lumen unless limited by working distance. A step size of 0.75 µm was used. The images were deconvoluted with the Lightning program from Leica. The image stacks were then imported into FIJI where sections were used to make a maximum Z projection of the hemisphere above the Geltrex hydrogel. A pipeline in CellProfiler 2 was then used to accentuate the ZO1 labeled tight junctions. ∼30 apical surface areas were measured by hand in FIJI for each maximum projection avoiding edges. We sought to measure all the surface areas in the center of the ZO1 labelled lumen to avoid size bias or surfaces orthogonal to the imaging plane.

## RESULTS

### High content imaging pipeline for SOSRS NTD screening

We have previously reported a protocol for generating self-organizing single rosette spheroids with manual imaging and analysis^37^. To dramatically increase throughput and decrease human bias, we have developed an entirely automated pipeline for imaging and analysis. First, we generate the neuroepithelial monolayer by 4 inhibitor treatment of iPS cells for 4 days. At this time, we cut the monolayer into identically size fragments and add the fragment suspension into a 96-well plate coated with Geltrex hydrogels (Figure 1a). After 48 hours of exposure, the cultures are paraformaldehyde fixed and stained. We then utilize a Yokogawa CV8000 automated dual microlens confocal to generate 10x images of each well of the 96-well plate containing on average 20-40 SOSRS per well (Figure 1b). Confocal optical sections are compressed into a maximum projection image (Figure 1c). Utilizing a CellProfiler pipeline, each SOSRS is identified from these images as well as the central lumen labeled with either ZO1-EGFP fusion protein, ZO1 antibody, or phalloidin for labeling f-actin (Figure 1d). Over 400 features including shape, fluorescence intensity, radial distribution, and area metrics are extracted from the images (Figure 1e). The SOSRS datasets are then filtered for individual lumens and size (7,800-31,415 µm^2^). We use these filters to exclude SOSRS that result from the fusion of multiple monolayer fragments as shown in our previous publication^37^. To identify distinguishing features for particular treatments or genetic variants, we utilize the random forest predictive model followed by inspecting the individual feature importance. For pharmaceutical treatments, linear regression random forest decreases the risk of false feature identification. Distinguishing features are then validated by univariate comparisons and statistical testing.

**Figure 1.**
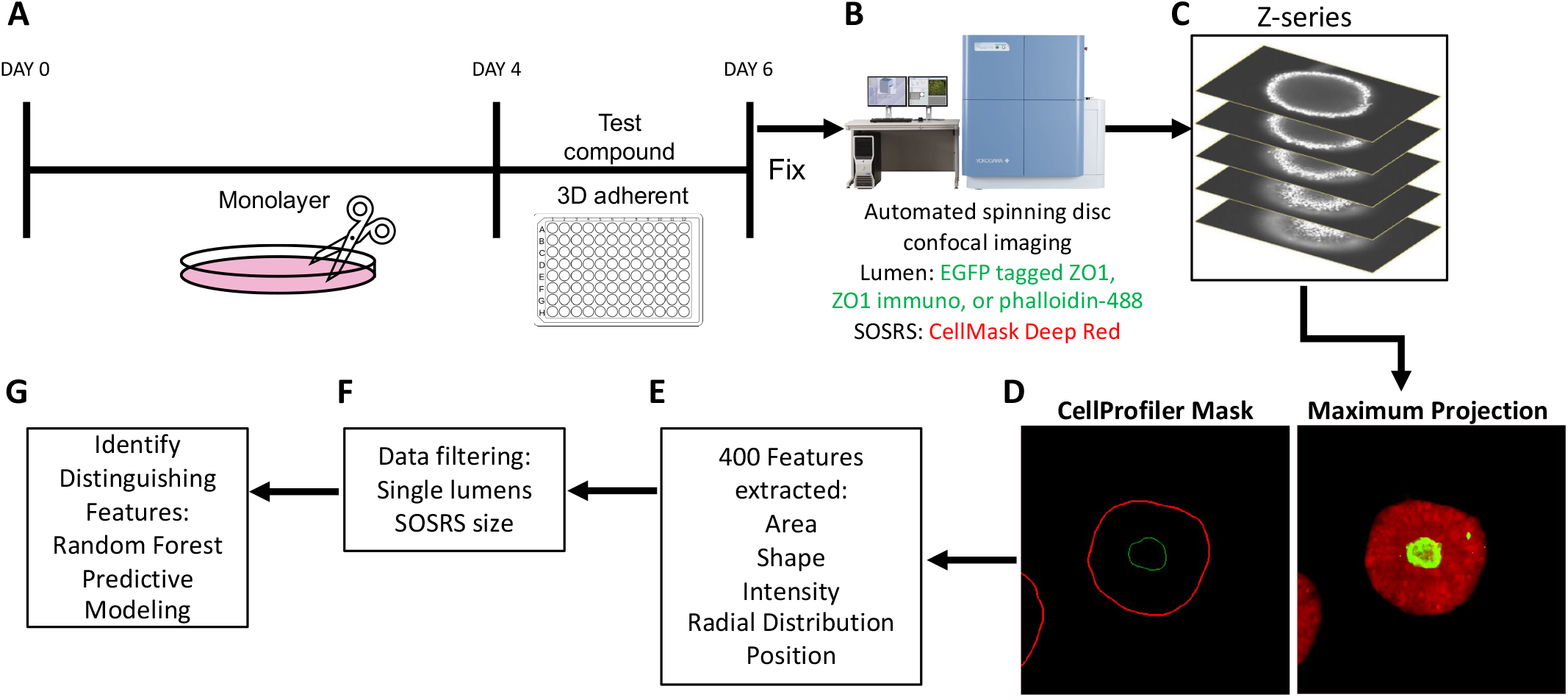
Automated platform for SOSRS NTD phenotyping. (**a**) Schematic of treatment paradigm of drug treatment during SOSRS neurulation. Monolayer neuroepithelial cultures were differentiated from iPS cells over 4 days. The monolayer was then cut with the StemPro EZ passage tool into reproducible fragments. These fragments were moved to a 96-well plate containing Geltrex and test compounds. Cells were cultured for 48 hours followed by paraformaldehyde fixation and counterstaining. (**b**) SOSRS were imaged for the constitutively tagged ZO1-EGFP and HCS CellMask Deep Red counterstain using the Yokogawa CellVoyager 8000. (**c**) 7 optical sections through the center of the SOSRS were generated. (**d**) The optical sections were used to generate a maximum projection in the far-red and green channels. CellProfiler was used to identify the margins for each channel for each SOSRS. (**e**) CellProfiler extracted over 400 features from the SOSRS in the two channels including area, shape, intensity, radial distribution, and position metrics. (**f**) Data was filtered for individual lumens and SOSRS area (7800-31415 µm2). (**g**) Random forest model is used to identify distinguishing features based on treatment.

### VPA dose response shows SOSRS lumen size as defining feature

VPA, a well-known cause of NTDs in mice and humans, previously gave consistent expansion of the lumen area normalized to the entire SOSRS area at 200 and 400 µM doses^37^. Here we performed a larger dose-response curve from 50 to 800 µM using our newly established pipeline. Our previously engineered feature, normalized lumen area, showed consistent increases across the entire dose-response curve (Fig. 2a,b). With the increased statistical power from the large number of SOSRS imaged by the automated confocal, we see a significant difference between any two concentrations by one-way ANOVA with a multiple comparison post hoc test. The same relationship can be seen from our original manually collected data across 3 independent experiments in Figure S1a. Using a linear regression based bootstrap random forest predictive model (Figure S1b) with the entire dataset (>400 extracted features) we found a high degree of predictiveness (R^2^ = 0.945) (Figure S1c). This model held true for the standard 5-fold cross validation (R^2^ = 0.811) (Figure S1d). The normalized lumen area (area ratio) was the most instructive feature for identifying the group with 30% of the overall impact (Figure S1e). All other top instructive features are almost entirely radial distribution of the ZO1-EGFP fluorescence-based features collectively making up 41.5% of the overall impact. The radial distribution proved to be useful in cases where the lumens were extremely large or had dysmorphic shapes as it did not require outlining the lumen, but instead relied on the fluorescent label of the lumen within the SOSRS mask. The radial distribution utilizes 6 concentric shapes within the SOSRS with identical diameter and the amount of EGFP signal is quantified in each of these rings. We normalized the data at each concentration to the most intense ring to account for the reduced signal intensity as the lumen expands. We then generated a three-color heatmap for plots of individual concentration averages and the entire dose response curve (Figure 2c,d). The gradual shift in fluorescence from primarily the inner three rings to the outer three rings is apparent.

**Figure 2.**
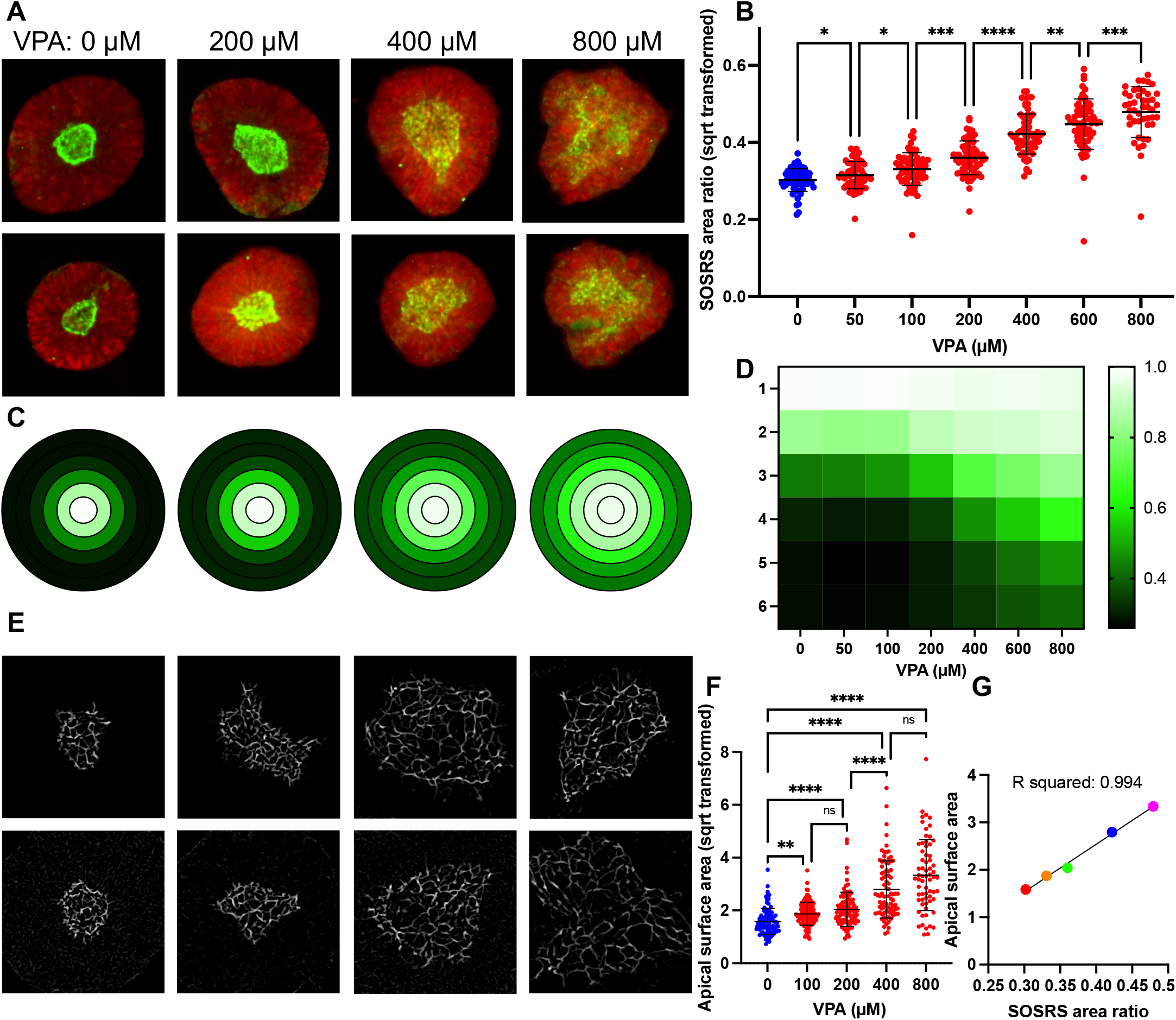
VPA causes a dose-response increase in SOSRS lumen size due to reduced apical constriction. (**a**) Example images of automated microscopy images of SOSRS treated with 0, 200, 600, and 1000 µM VPA. Green is ZO1-EGFP, and red is HCS CellMask Deep Red. (**b**) SOSRS lumen size (normalized to total SOSRS area and square root transformed) is plotted for VPA doses from 50 to 1000 µM. Each dot is an individual SOSRS. n = 63, 54, 66, 64, 68, 69, 45, and 32, respectively. Data from replicate independent experiments collected by manual imaging and analysis are presented in Figure S1a. A one-way ANOVA was used with FDR corrected post hoc test between groups. (**c**) Mean fraction of radial distribution plots (maximum normalized) for ZO1-EGFP fluorescence are depicted for the 4 concentrations in A. Each concentric ring represents the 6 concentric rings of equal thickness generated from each individual SOSRS and averaged for each group. Legend in D. (**d**) Heatmap of mean fraction radial distribution of ZO1-EGFP fluorescence across the entire VPA dose-response curve. Intensity is shown by 3 color heatmap with white as the highest intensity, black as the lowest, and green as intermediate. (**e**) Example 100x confocal images of the ZO1-EGFP labeled lumens with increasing VPA concentrations (0, 200, 400, 800 µM, respectively). (**f**) The size of individual cell apical surfaces as measured by the space in between the tight-junction is plotted. Kruskal-Wallis statistical test was performed with Dunn’s multiple comparison post hoc between groups. (**g**) XY scatter plot comparing the lumen max projection area to cell apical surface area means at each concentration of VPA. Colors are for increasing doses of VPA (red = 0, orange = 100, green = 200, blue = 400, and magenta = 800 µM VPA). Line and R2 value for linear regression analysis. Bars are mean and standard deviation. * p < 0.05, ** p < 0.01, *** p < 0.001, **** p < 0.0001.

### Lumen enlargement is directly related to reduced apical constriction

Since apical constriction is a necessary mechanism of neurulation and inhibitors of apical constriction lead to NTDs in mouse models and increased lumen size in our SOSRS model^37^, we decided to measure the apical surface areas of SOSRS treated with the dose curve of VPA. We utilized ZO1 immunostaining to increase the fluorescent signal for high magnification confocal microscopy (100x). This tight junction marker outlines each cell at the apical surface, and these outlines were measured to determine the area of each apical surface. We observed a dramatic increase in apical surface areas with increasing doses of valproic acid (Figure 2e). These results were significant compared with vehicle starting at 100 µM VPA and increasing up to 800 µM (Figure 2f). We then compared the relationship between the lumen size and apical surface area size by plotting the means of each at 0, 100, 200, 400, and 800 µM VPA on an XY scatter plot (Figure 2g). Linear regression analysis showed a striking level of correlation between these two measurements with an R^2^ of 0.994. Therefore, we believe measurement of the lumen area is a reasonable proxy measurement for cell apical surface areas in a given SOSRS brain organoid. This is expected because 3D lumen surface area is the sum of all the apical surface areas; the lumen must have a larger size to accommodate the larger individual surfaces.

### GSK3β and HDAC inhibition mimic VPA findings but no evidence for folic acid or oxidative stress involvement

With the clear phenotypic characteristics of the VPA-treated SOSRS, we attempted to find compounds that would ameliorate or mimic the effects of VPA on the lumen size. First, we attempted to block the effects of VPA with folic acid supplementation, the most effective prophylactic treatment for NTDs and a possible VPA target^13^. We also tested whether aminopterin, a potent inhibitor of the folic acid pathway, could mimic the effects of VPA. We found no effect of 10 µM folic acid on the normalized lumen size at 0, 200, or 400 µM VPA (Figure 3a). Conversely, we tested whether blocking folic acid synthesis using the dihydrofolate reductase inhibitor, aminopterin, could mimic the effects of VPA. Aminopterin exposure did not affect the lumen size (Figure 3b); thus, the evidence suggests that VPA does not alter lumen structure and apical constriction via inhibition of the folic acid pathway. One caveat is that our organoid cultures may be resistant to aminopterin due to an abundance of hypoxanthine and thymidine in the growth medium that can be used to generate purines through the salvage pathway without needing folic acid metabolism. Amino acid synthesize from folic acid pathway precursors are also abundant (i.e., glycine, serine, methionine). Thus, the major pathogenic mechanisms of folic acid deficiency are likely avoided.

**Figure 3.**
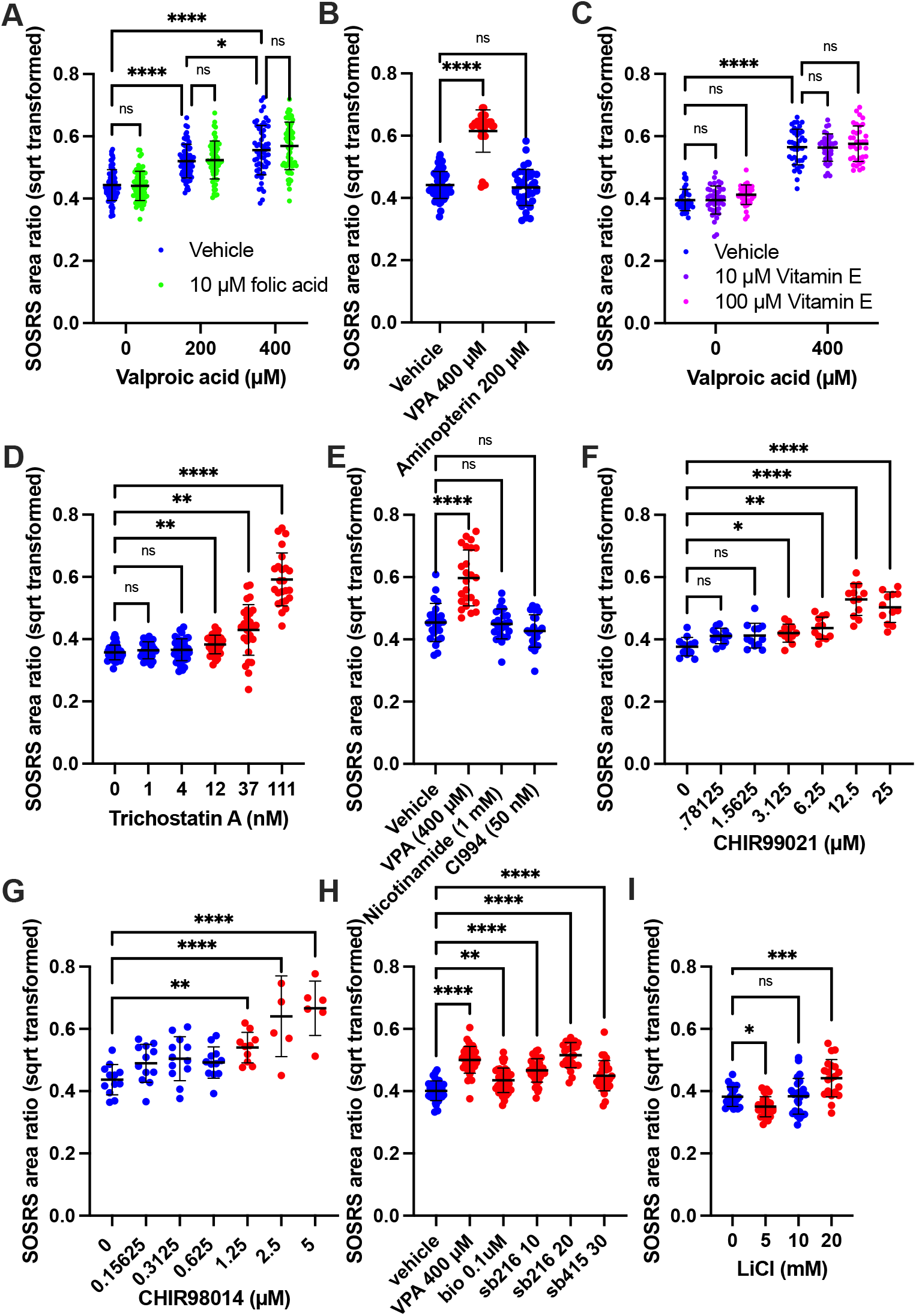
HDAC and GSK3β inhibitors mimic VPA-induced reduction of apical constriction. (**a-h**) SOSRS were treated with various compounds with or without valproic acid, and the lumen area normalized to the total SOSRS area and square root transformed is plotted for each set of experiments. Each dot is an individual SOSRS. The number of total SOSRS and total independent experiments are listed below. Analysis was performed by manual microscopy and blinded semi-automated analysis. (**a**) SOSRS were treated with vehicle, 200 or 400 µM valproic acid with or without 10 µM folic acid in addition to the basal media concentration (7.5 µM). n = 71, 71, 66, 63, 56, and 66 respectively across 3 independent experiments. (**b**) SOSRS were treated with vehicle, 400 µM VPA, or 200 µM aminopterin (folic acid pathway [DHFR] inhibitor). n = 55, 34, and 37, respectively across 2 independent experiments. (**c**) SOSRS were treated with vehicle or 400 µM VPA with the addition of 0, 10, or 100 µM vitamin E (antioxidant). n = 40, 40, 36, 40, 36, and 36, respectively across 2 independent experiments. (**d**) SOSRS were treated with a dose curve of the class I/II HDAC inhibitor, trichostatin A. n = 40, 28, 39, 38, 28, and 24, respectively across 3 independent experiments. (**e**) SOSRS were treated with HDAC inhibitors nicotinamide (1 mM) and CI994 (50 nM) and compared to 400 µM VPA. n = 24 for each group across 2 independent experiments. (**f**) SOSRS were treated with a log2 dose curve of the GSK3β inhibitor CHIR99021. n = 12 for each group for 1 experiment. (**g**) SOSRS were treated with a log2 dose curve of the GSK3β inhibitor CHIR98014. n = 12, 12, 12, 12, 10, 6, and 6, respectively for 1 experiment. (**h**) SOSRS were treated with VPA and compared with other GSK3β inhibitors BIO, SB216763, and SB415286. n = 36, 36, 36, 36, 24, and 29, respectively across 3 independent experiments. (**i**) SOSRS were treated with 0, 5, 10, and 20 mM lithium chloride. n = 24, 24, 24, and 20, respectively across 2 independent experiments. Bars are mean and standard deviation. Statistical analyses were performed by two-way ANOVA with Tukey post-hoc test (**a,c**) and one-way ANOVA with Dunnett’s multiple comparisons test (**b,d-i**). * p < 0.05, ** p < 0.01, *** p < 0.001, **** p < 0.0001.

To test the hypothesis that VPA caused NTDs by increasing oxidative stress^16^, we added antioxidants, vitamin E or resveratrol, during VPA exposure. Again, there was no effect of either 10 or 100 µM vitamin E or 10 µM resveratrol on the lumen size for vehicle or 400 µM VPA treatment (Figure 3c and data not shown). Again, these data suggest that increased oxidative stress is not the mechanism by which VPA causes decreased apical constriction. The most well-known action of VPA is HDAC inhibition^14^. Treatment with HDAC I and HDAC III inhibitors, nicotinamide and CI994 respectively, did not affect lumen size, but trichostatin A (TSA), an HDAC I/II inhibitor as is VPA, caused a similar dose-response increase in lumen size (Figure 3d,e). High magnification microscopy also showed that 100 nM TSA causes dramatic enlargement of apical surfaces areas compared with vehicle and to the same extent as 800 µM VPA (Figure S2a,b). VPA is also known to inhibit GSK3β^15^. Inhibitors of GSK3β including CHIR99021, CHIR98014, SB216763, SB415286, BIO, and lithium all caused increased lumen size (Figure 3f-i). Exemplary images showing the increased lumen size are shown in Figure S2c Therefore, both HDAC and GSK3β inhibition decrease apical constriction, and one or both are the likely mechanisms by which VPA causes NTDs.

### RNA sequencing results suggest VPA inhibits HDAC but not GSK3β at concentrations causing NTD phenotypes

Since HDACs and GSK3β signaling both have large transcriptomic effects, we characterized the transcriptomic changes caused by concentrations of VPA (at 200 and 400 µM) that inhibited apical constriction. We also compared the effects of a specific GSK3β inhibitor, CHIR99021 (10 µM). Four independent experimental sample sets were generated by extracting RNA from SOSRS after 48 hours of drug exposure. Principal component analysis showed excellent clustering by treatment in PC2 while a single experimental replicate was highly divergent in PC1 (Figure 4a). At 200 µM, VPA exposed SOSRS had 800 differentially expressed genes (DEG = > 50% change and a padj of < 0.05) and 1833 at 400 µM with 783 shared DEGs. CHIR99021 exposure had 699 DEGs with only 70 and 170 DEGs shared with 200 and 400 µM VPA, respectively (Figure 4b). Despite massively altering the transcriptome, the most striking effect of VPA was an increase in the expression of neuronal genes. The top cellular compartment GO terms for increased VPA DEGs were nearly all neuronal including “axon”, “neuronal cell body”, “postsynaptic membrane”, “synapse”, and “dendrite” (Figure 4c), and the top increased DEG for both 200 and 400 µM was *STMN2*, a quintessential transcriptional marker of neurons (Figure 4d,e). This effect on neuronal genes has been previously shown in neural progenitors due to increased acetylation of H4 which associates with the proneuronal *Ngn1* promoter^41^. Inhibition of GSK3β via CHIR99021 had many different top transcripts from VPA although *STMN2* was still increased but to a lesser extent (Figure 4f). Inhibition of GSK3β mimics the effects of WNT signaling and should result in increased expression of TCF/LEF target transcripts. Indeed, targets *TCF7, DKK1, AXIN2,* and *CCND2* were all significantly increased with CHIR exposure (Figure 4g); however, these transcripts were not significantly affected by 200 or 400 µM VPA. Furthermore, GO analysis of DEGs showed that WNT was a significantly affected pathway by CHIR but not VPA (Figure 4h-j, yellow circle indicates the WNT pathway). These data show that VPA is not inhibiting GSK3β activity at these concentrations, and thus GSK3β is not the cause of the increased lumen area in VPA treated SOSRS. More likely, the massive effect on the transcriptome due to HDAC inhibition is causing the structural changes. Lastly, genes important for neurulation and associated with NTDs (e.g., *SHROOM3*, *NUAK2*, *VANGL2, CELSR2*) were all decreased by VPA and CHIR99021 exposure (Figure 4k). Data for all expressed genes comparing between treatment and vehicle controls is displayed in Tables S4-6.

**Figure 4.**
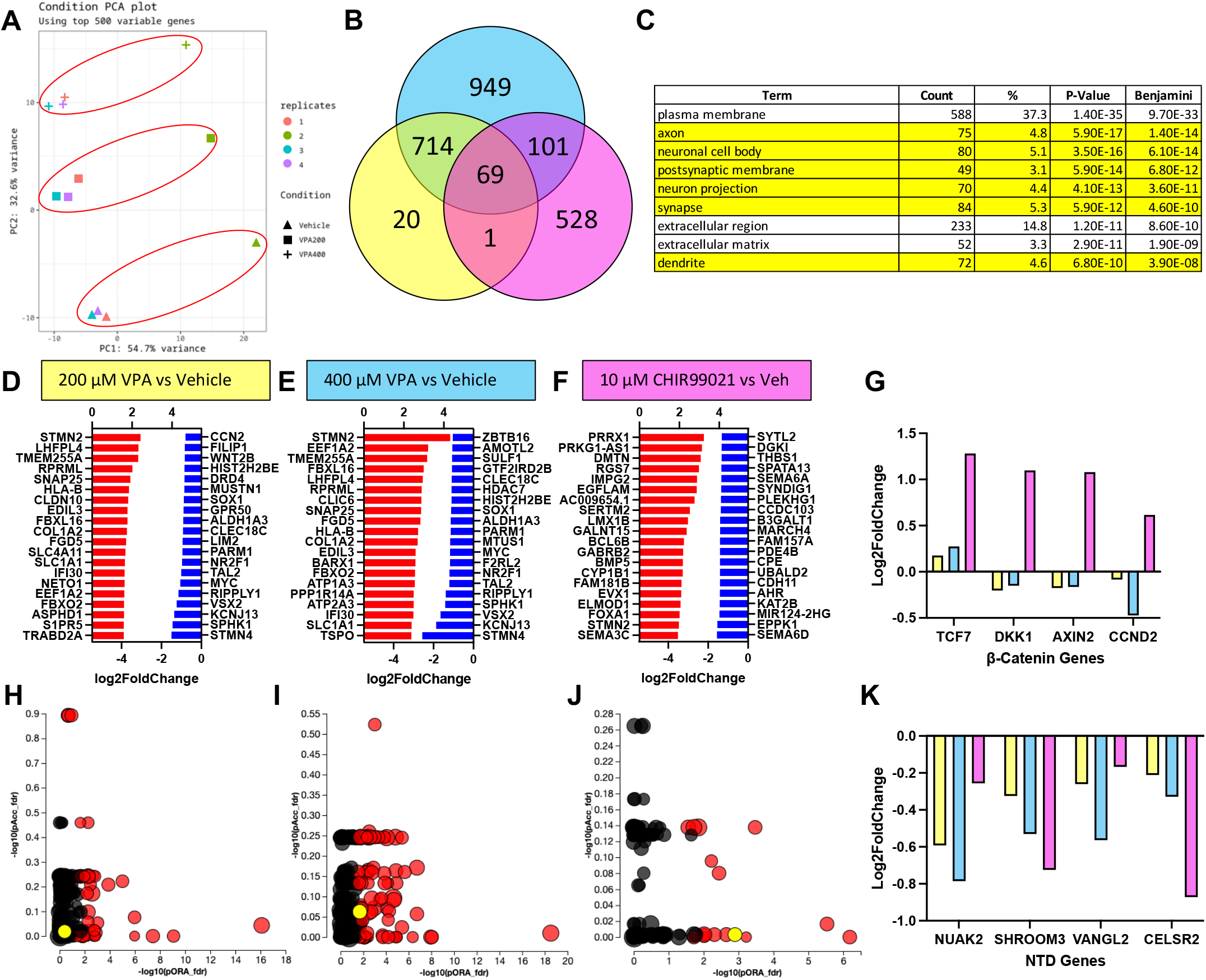
VPA causes transcriptomic signatures of neural differentiation but not GSK3β inhibition. (**a**) Principal component analysis of VPA (200 and 400 µM) and vehicle-treated SOSRS mRNA sequencing analysis for 4 independent replicates. Red circles indicate treatment groupings. (**b**) Venn diagram of differentially expressed genes (DEGs) for 200, 400 µM VPA and 10 µM CHIR99021 compared with vehicle control in SOSRS. Overlapping areas indicate the number of shared DEGs between treatment groups. (**c**) Gene ontology analysis of upregulated DEGs for 400 µM VPA treatment based on protein subcellular localization. Of the 9 top terms, 6 are neuron-specific (i.e., axon, dendrite, synapse). (**d-f**) The top 20 upregulated and downregulated transcripts for each treatment: (**d**) 200 µM VPA, (**e**) 400 µM VPA, and (**f**) 10 µM CHIR9901. (**g**) Log2 fold change in transcript levels for 4 canonical Wnt/β-catenin pathway regulated genes for each treatment. Only the GSK3β inhibitor CHIR99021 significantly increased the transcript levels for each of these genes. (**h-j**) Scatter plots of pathway GO terms by two significance scores (pORA: p-value for over-representation analysis, pAcc: p-value from total perturbation accumulation for the pathway) corrected for false discovery rate. A data table for significant terms is listed in Table S3. The yellow circle in each plot is the Wnt pathway. (**h**) 200 µM VPA, (I) 400 µM VPA, and (J) 10 µM CHIR99021. (**k**) Log2 fold change in transcript levels for 4 NTD-associated genes for each treatment. Both VPA and CHIR99021 decrease transcript levels for each of these genes.

### SHROOM3 knockout results in enlarged, dysmorphic lumens and reduced apical constriction

To further investigate the utility of our NTD model system, we sought to generate a genetic model of NTDs. *SHROOM3* knockout mice have exencephaly, and several human anencephaly pregnancies with *SHROOM3* variants have been reported^22^. We, therefore, utilized our previously developed methodology for generating knockout iPSC lines by simultaneous reprogramming and CRISPR indel formation to generate a *SHROOM3* knockout iPSC line and isogenic control^39^. We used a gRNA in the same exon as a human anencephaly pregnancy with a premature stop codon mutation (Figure 5a)^22^. We generated a line with compound heterozygous frameshift mutations via −1/−14 bp indels at the CRISPR/Cas9 cut site as well as an isogenic control without indel formation (Figure 5b-d). Utilizing quantitative reverse transcriptase PCR, we found that *SHROOM3* transcript levels peaked on day 4 of differentiation while *PAX6* transcript also peaked (Figure S3a). This fits with prior studies that find *SHROOM3* expression is dependent on *PAX6* expression^42^. qRT-PCR did not show any difference between the isogenic control and our engineered knockout line, indicating a lack of nonsense-mediated decay (Figure S3b). Immunoblot analysis had too many non-specific bands to confirm knockdown (data not shown); however, using immunocytochemistry for *SHROOM3* on whole mount SOSRS, we see a ring of apical SHROOM3 expression in the isogenic control that is absent in the knockout line (Figure S3c,d). Furthermore, we see a dramatic structural change in the SOSRS lumen morphology. Instead of a small circular lumen as seen in the isogenic controls (Figure 5e,j, Figure S3e), we observed massively enlarged lumens with lobes protruding into the lumen space (Figure 5f,k, Figure S3f). Both f-actin staining via a phalloidin-fluorophore conjugate (Figure 5e,f, Figure S3f) and ZO1 immunostaining (Figure 5j,k) show these clear phenotypes, but the f-actin staining also shows basal expression only in the *SHROOM3* knockout (Figure 5f arrowheads) while ZO1 is only on the apical lumen surface in either line (Figure 5j,k). We even see the occasional *SHROOM3-*KO SOSRS with a large opening at the top of the lumen suggesting incomplete closure (Figure 5k). Using our high throughput imaging and feature identification, we found that, radial distribution of either marker, lumen solidity, and f-actin staining on the SOSRS edge are excellent distinguishing features (Figure 5h,i,m). Except for the basal f-actin, these features were identified for both ZO1 and f-actin stained SOSRS (Figure S4).

**Figure 5.**
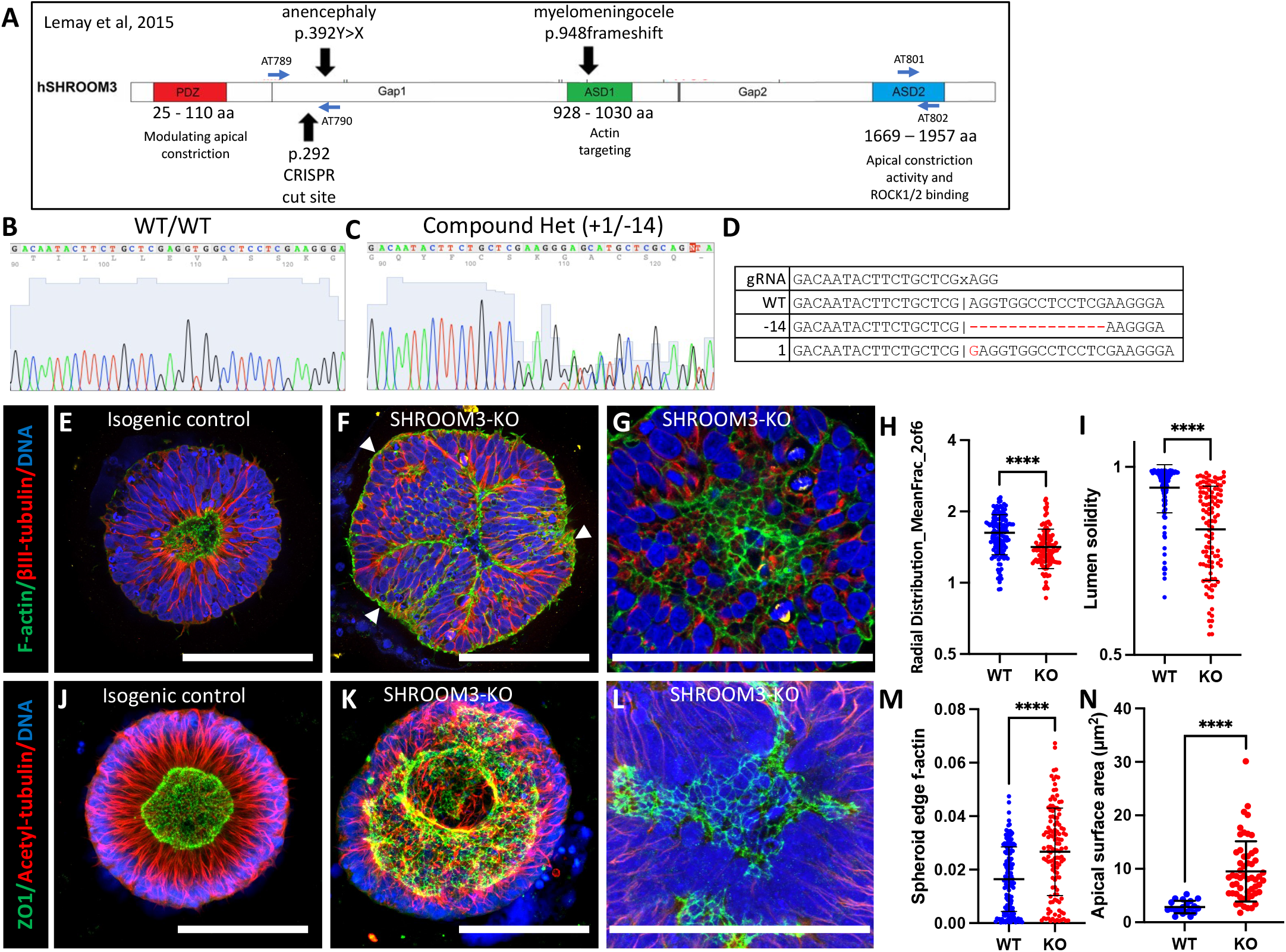
SHROOM3 knockout SOSRS have enlarged dysmorphic lumens, basal f-actin, and impaired apical constriction. (**a**) Schematic of human SHROOM3 gene with important locations such as known human NTD variants, CRISPR cut site, and PCR primer locations. (**b,c**) Sanger sequencing chromatographs for an isogenic control and compound heterozygous frameshift indel line using PCR primers AT789 and AT790. (**d**) The gRNA and indel sequences are listed. The “x” indicates the Cas9 cut site in the sequence. (**e-g**) Confocal images of SOSRS immunostained for β-III-tubulin and stained with f-actin dye, phalloidin-Alexa488. (**e**) Isogenic control has typical small round, central lumen with radial tubulin structure. (**f**) SHROOM3-KO has enlarged, dysmorphic lumen and basally localized f-actin (arrowheads). (**g**) Digital magnification of maximum Z-projection of one side of the lumen to show enlarged apical surface areas. (**h**) Knockout cells have reduced f-actin fluorescence mean fraction in the ring 2 of 6 compared with controls. (**i**) Knockout cells have reduced lumen solidity compared with controls. (**j-l**) Confocal images of SOSRS immunostained for acetyl-tubulin and ZO1. (**j**) Isogenic control has typical small round, central lumen with radial tubulin structure. (**k**) SHROOM3-KO has enlarged, dysmorphic lumen with large opening (max project is of the entire lumen Z-stack). (**l**) Digital magnification of maximum Z-projection of one side of the lumen to show enlarged apical surface areas. (**m**) Knockout cells have increased f-actin fluorescence on the SOSRS outer edge compared with controls. (**n**) Knockout cells have greater individual cell apical surface areas than isogenic controls. Each dot in **h**, **i**, **m** represents measurements for an individual SOSRS while in **n** they represent individual apical cell surface areas. n = 130 for WT and 117 for KO for **h,i,m**. n = 20 for WT and 55 for KO in **n**. Bars are mean and standard deviation. Statistical analysis performed with Mann-Whitney test. Scale bars = 100 µm. * p < 0.05, ** p < 0.01, *** p < 0.001, **** p < 0.0001.

Again, our previous work indicates that lumen enlargement is due to reduced apical constriction. To test this hypothesis for *SHROOM3* knockout, we performed high-resolution microscopy and area measurements for apical surface areas as defined by the tight-junction marker, ZO1. The size changed from 2.9 ± 1.2 µm^2^ (n = 20) for the isogenic control to 9.5 ± 5.6 µm^2^ for the knockout line (Figure 5n). The large apical surface areas can be seen by either f-actin or ZO1 immunostaining (Figure 5g,l, respectively). These data fit with the literature that *SHROOM3* is necessary for polarized localization of apical constriction machinery and subsequent constriction. Therefore, reduced apical constriction is expected as is the increased basal f-actin. Both have been seen as a consequence of *SHROOM3* loss of function in a mouse model^42^.

### Mosaic SHROOM3 knockout shows gene-dose response and possible non-cell autonomous effects

It is theorized that NTDs can be caused not only by germline mutations but also somatic mutations in the neural tube resulting in mosaic loss-of-function of important proteins. One recent publication found that knocking out *VANGL2* in only 16% of the cells in the neural tube of a developing mouse embryo was sufficient to produce spina bifida^26^. Since we have both *SHROOM3*-KO and isogenic control lines, we sought to investigate this hypothesis by mixing the two genotypes at 5 different ratios (% *SHROOM3*-KO: 0, 25, 50, 75, and 100). We found the same features (radial ZO1 and f-actin distribution, lumen solidity, and SOSRS edge f-actin) distinguished the 0 and 100% with the mixed cultures showing a clear intermediate dose response for each measurement (Figure 6a-c). Interestingly, 25% KO was significantly different from 0% for each measurement, but 75% and 100% were indistinguishable from one another for each measurement. This may suggest a non-cell autonomous effect of *SHROOM3* knockout as suggested in *Xenopus* studies^43^. Similar to the VPA dose-response, we graphed the gene-dose effect on radial distribution of ZO1 and f-actin using a heatmap (Figure 6d-h). These data clearly show the lumen size increases in direct relationship with the increased percentage of *SHROOM3*-KO cells.

**Figure 6.**
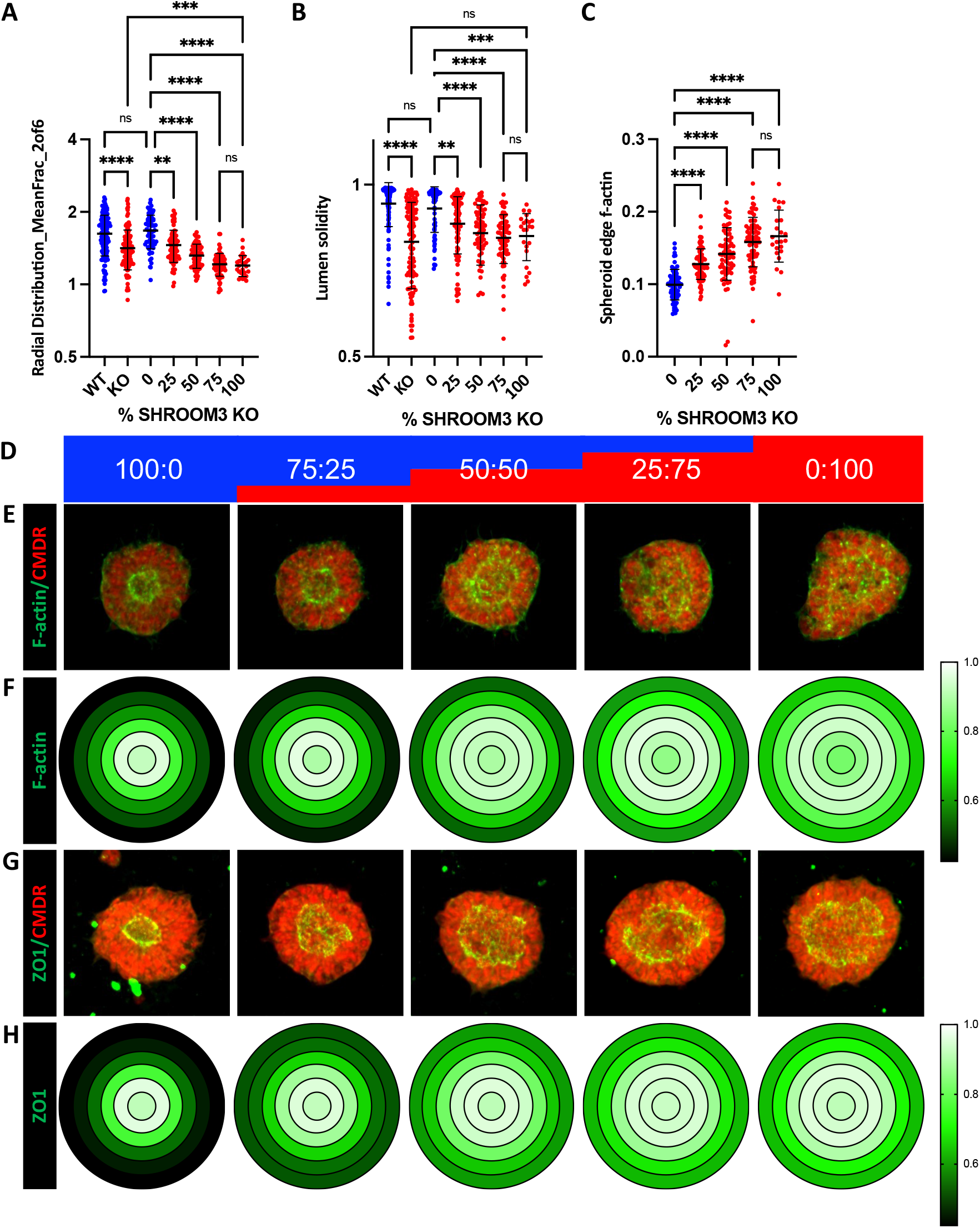
Mosaic mixing of WT and KO SHROOM3 lines results in gene-dose response and suggests non-cell autonomous effects. (**a,b**) Comparing distinguishing features from figure 5 dataset and 5 different percentage doses of SHROOM3 KO cells (0, 25, 50, 75, and 100 %, remainder of cells are WT). (**a**) Data for f-actin mean fraction of fluorescence radial distribution in ring 2. (**b**) Lumen solidity for each group. (**c**) The mean intensity of f-actin on the SOSRS basal edge was measured and a similar gene dose response was seen. n = 55, 68, 58, 59, and 22, respectively across 2 independent experiments. Comparisons were not made with the Figure 5 dataset due to differences in overall fluorescence signal because of different phallodin-Alexa488 vendors. (**d**) A schematic of the mixing of SHROOM3-KO and isogenic WT cells by percent. (**e**) Automated confocal maximum projections for exemplary SOSRS at each of the 5 mixes labelled for f-actin with phalloidin-Alexa488 and HCS CellMask Deep Red. (**f**) Average maximum normalized radial distribution of mean fraction of f-actin in each of the 6 concentric ring bins. Intensity is shown by 3 color heatmap with white as the highest, black as the lowest, and green as intermediate. (**g**) Automated confocal maximum projections for exemplary SOSRS at each of the 5 mixes immunolabelled for ZO1 and HCS CellMask Deep Red. (**h**) Average maximum normalized radial distribution of mean fraction of ZO1 in each of the 6 concentric ring bins. Intensity is shown by 3 color heatmap with white as the highest intensity, black as the lowest, and green as intermediate. Bars are mean and standard deviation. Statistical analysis performed with Kruskal-Wallis with Dunn’s multiple comparisons test. * p < 0.05, ** p < 0.01, *** p < 0.001, **** p < 0.0001.

## DISCUSSION

Our results show that our model system is useful for rapid screening of pharmaceuticals and genetic variants for NTD-like structural phenotypes. We have shown dose-responsive increases in lumen size for VPA that correlates with decreased apical constriction. We also show that VPA has this effect primarily through HDAC inhibition. In this report, we also generated a genetic anencephaly model by knocking out *SHROOM3,* which resulted in increased lumen size and reduced apical constriction, similar to the effects of VPA. Additionally, the *SHROOM3* knockout also altered polarity of filamentous actin and the circularity of the lumen structure. The effects of *SHROOM3* knockout were also shown to be gene-dose responsive in our mosaic mixing experiments, and these results indicated a possible non-cell-autonomous effect and that an open NTD could result when only a minority of cells in the neural tube contain a pathogenic variant. Taken together, these data suggest that reduced apical constriction is a shared mechanism for these two quintessential causes of NTDs. Furthermore, these results along with the robust high-throughput platform we have developed, and the statistical power of the model system suggest that our system would perform well for screening for novel pharmaceutical or genetic causes of NTDs.

Inhibitors of apical constriction have been shown previously in our system and in rodent embryo models to result in enlarged dysmorphic lumens and NTDs, respectively ^37, 44^. *SHROOM3* has been shown to be necessary and sufficient to induce apical constriction in cell culture and embryo models^45^, but the mechanism by which VPA leads to NTDs has been less clear. While our data indicate that VPA causes NTDs via HDAC inhibition, which of the 800+ genes with altered expression and what functions are impinged are not clear from transcriptomic data alone. Our structural phenotyping clearly shows that apical constriction is reduced by VPA. VPA also led to a decrease in the expression of 4-NTD related genes, but further study is needed to understand which transcriptional changes lead to outcome of reduced apical constriction. Our data differ dramatically from one recently published study which found 1 µM VPA was sufficient to disrupt single rosette formation by inhibiting apical f-actin and that this effect could be blocked by the addition of 10 µM folic acid^36^. Our study shows 50 µM VPA is needed to see clear evidence of lumen enlargement and that 10 µM folic acid had no effect on the structural changes seen with 200 or 400 µM VPA. One key difference is that the former published study treated cultures with VPA starting at day 0 of neural induction, while our study added VPA when neurulation was induced on day 4. Our transcriptomic data showed a modest but dose-response decrease in *SHROOM3* expression with VPA treatment. The loss of apically polarized f-actin but not ZO1 is strikingly similar to our *SHROOM3* knockout findings^36^. Thus, early administration of VPA may inhibit *SHROOM3* expression resulting in mislocalized f-actin. However, the differences in folic acid rescue between this study and ours would still not be explained unless folic acid somehow induces *SHROOM3* expression.

We also unexpectedly found that inhibition of GSK3β results in the same enlargement of the apical lumen and presumably decreased apical constriction. Our data also shows that lithium exposure, which has controversially been linked to NTDs, could be a cause at high concentrations due to inhibition of GSK3β. The mechanism is likely due to alterations in β-catenin signaling and downstream transcriptional alterations. Interestingly, CHIR99021 exposure in SOSRS also decreases *SHROOM3* expression similar to VPA. Therefore, inhibition of *SHROOM3* expression leading to reduced apical constriction could be a shared mechanism for VPA and GSK3β inhibitors and should be further explored, possibly by rescue experiments.

Like all model systems, the SOSRS model has limitations. While highly amenable to high-throughput techniques, there are still important aspects in whole embryos or multi-lineage systems that cannot be recapitulated. For instance, our current SOSRS methodology has no developmental axis patterning such as dorsal-ventral or rostral-caudal. The spheroids are all patterned to be dorsal forebrain^37^. This is vitally important to the study of spina bifida which occurs at the caudal rather than rostral extreme of the neural tube. Additionally, while many lines of data indicate spina bifida and anencephaly have many shared mechanisms, some causes seem to be specific to one or the other. Future work to develop a lumbar spinal cord patterned single-rosette CNS organoid would be useful in this regard.

In the future, the SOSRS model will need to be evaluated for predictive accuracy for a larger panel of known teratogenic and non-teratogenic compounds. If the predictive nature we see in the current study with VPA and *SHROOM3* continues for such panels, the SOSRS model may prove to be an ideal platform for identifying neuroteratogenic compounds during drug development or for existing compounds. These studies are vitally important for novel therapeutic screening and comparisons within drug classes known to carry NTD risk, such as antiseizure medications and possibly HIV antiretroviral therapies ^10, 46^. Additionally, our current results indicate that possible genetic NTD risk variants can be assessed in this system, but this warrants further investigation of outcomes in a larger number of well-characterized NTD risk genes such as *VANGL2* and *SCRIB*^23, 47^.

## Supplementary Materials

Figure S1: SOSRS lumen area and radial distribution of ZO1-EGFP are distinguishing features of VPA treated SOSRS; Figure S2: HDAC and GSK3β inhibitors decrease apical constriction; Figure S3: Loss of apical SHROOM3 immunostaining confirms loss of functional protein; Figure S4: Radial distribution found as most instructive feature to determine SHROOM3 genotype percentage using either f-actin or ZO1 staining; Table S1: Primer and gRNA sequences; Table S2: Antibodies and dilutions; Table S3: Features extracted from ZO1-EGFP and CMDR labeled SOSRS.

## Author Contributions

Conceptualization, A.T.; methodology, A.T., J.W., and J.S.; software, J.H. and J.S.; validation, T.T., J.L., R.S., and A.T.; formal analysis, T.T, J.L., R.S., J.H., and A.T.; investigation, T.T., J.L., R.S., J.H., J.W., and A.T.; resources, J.S., J.P., and A.T.; data curation, J.H., J.S., and A.T.; writing—original draft preparation, A.T.; writing— review and editing, T.T., J.L., R.S., J.W., N.V., J.S., J.P., and A.T.; visualization, A.T.; supervision, N.V., J.P., and A.T.; project administration, A.T.; funding acquisition, N.V. and A.T. All authors have read and agreed to the published version of the manuscript.

## Institutional Review Board Statement

Not applicable

## Informed Consent Statement

Not applicable

## Acknowledgments

We acknowledge support from the Bioinformatics Core of the University of Michigan Medical School’s Biomedical Research Core Facilities (RRID:SCR_019168). We would also like to acknowledge the helpful input to this project from M. Elizabeth Ross, Richard Finnell, Louis Dang, and Kelly Gilmore.

