## Supplemental Figures S1-4 for "A Shared Pathogenic Mechanism for Valproic Acid and SHROOM3 Knockout in a Brain Organoid Model of Neural Tube Defects"

A

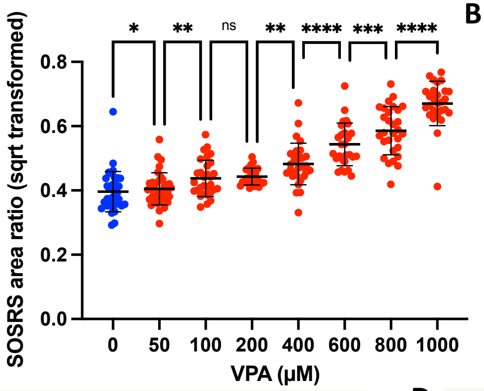


B

| Target | VPA Concentration |
|---|---|
| Number of Trees in the forest | 100 |
| Number of terms sampled per split | 247 |
| Training rows | 337 |
| Validation rows | 92 |
| Number of terms | 406 |
| Bootstrap samples | 337 |
| Training Rsquare | 0.945 |
| Validation Rsquare | 0.811 |

C D

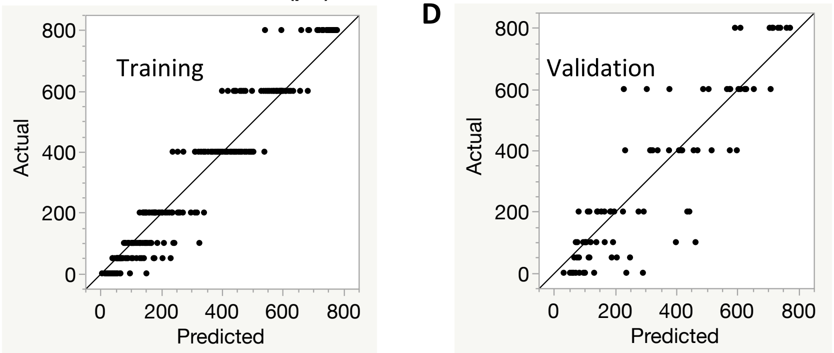


E

| Term | Number of Splits | SS | Portion |
|---|---|---|---|
| Area_Ratio | 65 | 4203854.01 | 0.2957 |
| spheroids_RadialDistribution_MeanFrac_GFP_4of6 | 26 | 2753430.53 | 0.1937 |
| spheroids_RadialDistribution_FracAtD_GFP_4of6 | 25 | 2688079.63 | 0.1891 |
| spheroids_RadialDistribution_ZernikeMagnitude_GFP_6_0 | 6 | 430509.521 | 0.0303 |
| spheroids_Intensity_MeanIntensityEdge_CMDR | 33 | 296548.113 | 0.0209 |
| spheroids_RadialDistribution_FracAtD_GFP_2of6 | 9 | 157367.336 | 0.0111 |
| GFP_Core_AreaShape_Solidity | 10 | 117949.747 | 0.0083 |
| spheroids_RadialDistribution_ZernikeMagnitude_GFP_7_1 | 6 | 112700.192 | 0.0079 |
| GFP_Core_AreaShape_Compactness | 9 | 93466.0962 | 0.0066 |
| GFP_Core_AreaShape_Zernike_0_0 | 9 | 87516.7566 | 0.0062 |
| spheroids_RadialDistribution_FracAtD_GFP_1of6 | 16 | 83750.4414 | 0.0059 |
| spheroids_RadialDistribution_MeanFrac_GFP_3of6 | 7 | 71758.7242 | 0.0050 |
| GFP_Core_AreaShape_Extent | 13 | 61462.9614 | 0.0043 |
| spheroids_RadialDistribution_RadialCV_GFP_2of6 | 9 | 59414.21 | 0.0042 |
| spheroids_RadialDistribution_FracAtD_GFP_3of6 | 7 | 58368.3585 | 0.0041 |

**Supplemental Figure 1.** SOSRS lumen area and radial distribution of ZO1-EGFP are distinguishing features of VPA treated SOSRS. (**a**) The normalized lumen area was quantified in SOSRS treated with a dose curve of VPA. Analysis was performed manually. N = 36, 36, 30, 26, 31, 29, 29, and 27, respectively across the 3 independent experiments. (**b**) Specific details and results for the random forest predictive model utilized with the automated data presented in Figure 2B. (**c**) Actual treatment concentration vs. predicted XY scatter-plot based on the bootstrap random forest model for the training data (80% of dataset). (**d**) Actual treatment concentration vs. predicted XY scatter-plot based on the bootstrap random forest model for the validation data (20% of dataset). (**e**) Top 15 most instructive features (of 407 total) for the prediction of VPA treatment concentration (from 50-800 µM). The portion of overall predicted value is the graphs and portion value out of a total of 1.

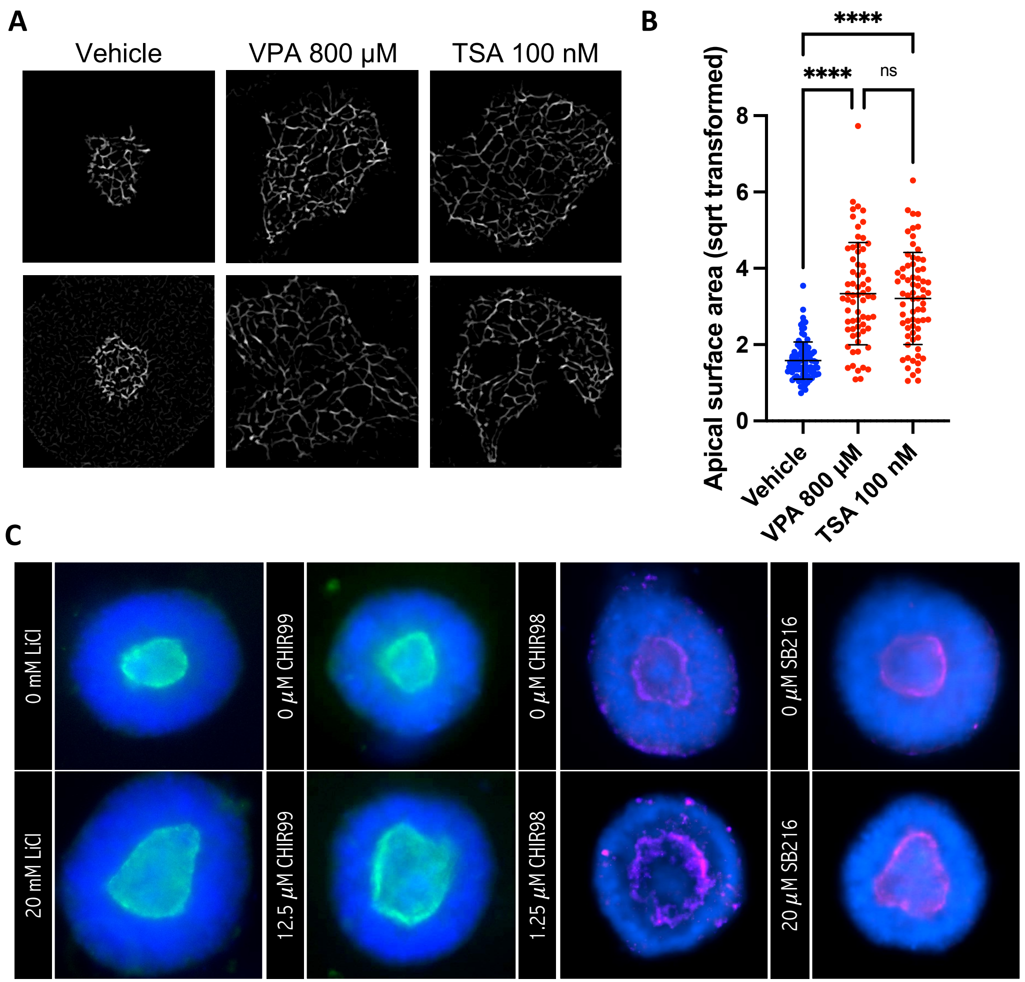


**Supplemental Figure 2.** HDAC and GSK3β inhibitors decrease apical constriction. (**a**) Apical cell surface areas were imaged by immunostaining for the tight-junction marker ZO-1 which outlines each cells apical surface. Two images for vehicle, 800 µM VPA, and 100 nM TSA are shown. (**b**) Individual surfaces were measured manually for n = 93 across 3 lumens, 66 across 2 lumens, and 69 across 2 lumens, respectively. (**c**) Exemplary manual images of SOSRS taken after labeling for DNA (bis-benzamide) and the apical lumen with either ZO1-EGFP (green) or ZO1 immunostaining (magenta). Immunostaining was necessary for some drugs due to fluorescence in the green channel interfering with ZO1-EGFP signal. Error bars are standard deviation. Statistical analysis performed with Kruskal-Wallis with Dunn's multiple comparisons post hoc. **** $p < 0.0001$.

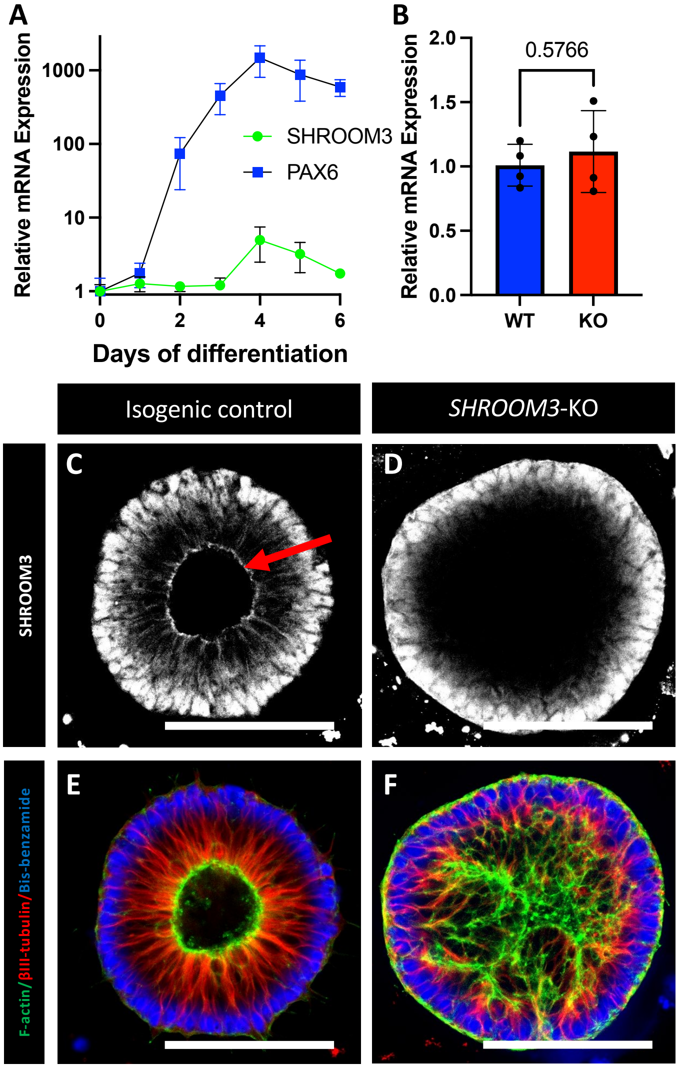


**Supplemental Figure 3.** Loss of apical SHROOM3 immunostaining confirms loss of functional protein. (**a**) Quantitative reverse transcriptase PCR was performed for *PAX6* and *SHROOM3* across 0-6 days of SOSRS differentiation. N = 2-4 independent differentiations for each data point. (**b**) The same qRT-PCR was performed on day 4 for both the *SHROOM3*-KO line and isogenic control. N = 2 samples for 2 independent experiments (4 total each group). Comparison was by unpaired t-test. Error bars are standard deviation. (**c**-**f**) Confocal micrographs of isogenic (**c**,**e**) and *SHROOM3*-KO SOSRS (**d**,**f**) stained for either SHROOM3 (**c**,**d**) or f-actin, beta-III-tubulin, and bis-benzamide (**e**,**f**). Scale bars are 100 µm.

**A**

| Target | F-actin | ZO1 |
|---|---|---|
| Number of Trees in the forest | 100 | 100 |
| Number of terms sampled per split | 268 | 262 |
| Training rows | 258 | 388 |
| Validation rows | 52 | 87 |
| Number of terms | 437 | 437 |
| Bootstrap samples | 258 | 388 |
| Training Rsquare | 0.900 | 0.863 |
| Validation Rsquare | 0.644 | 0.431 |

**B** F-actin

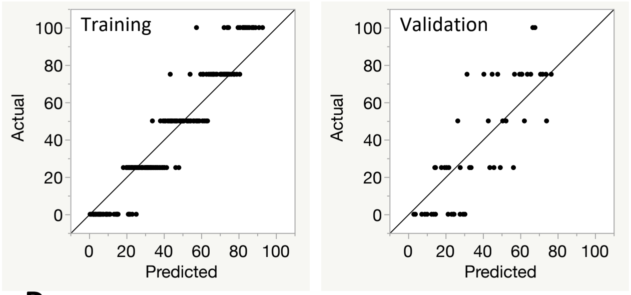


**C** ZO1

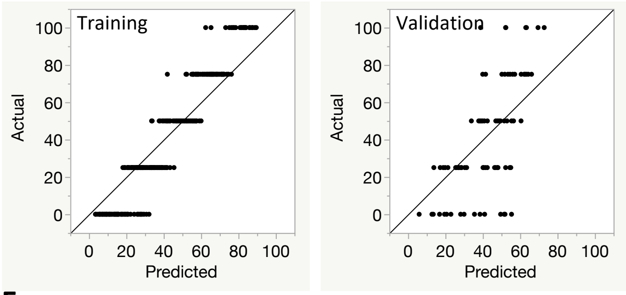


**D**

| Term | Splits | SS | | Portion |
|---|---|---|---|---|
| spheroids_RadialDistribution_MeanFrac_GFP_2of6 | 53 | 44453.7285 | | 0.2979 |
| spheroids_Intensity_LowerQuartileIntensity_GFP | 27 | 7260.78485 | | 0.0487 |
| spheroids_Intensity_MedianIntensity_GFP | 17 | 4506.9406 | | 0.0302 |
| spheroids_RadialDistribution_FracAtD_GFP_2of6 | 10 | 4317.22953 | | 0.0289 |
| spheroids_RadialDistribution_MeanFrac_GFP_4of6 | 13 | 3805.69547 | | 0.0255 |
| spheroids_RadialDistribution_MeanFrac_GFP_5of6 | 13 | 3632.0435 | | 0.0243 |
| GFP_Core_Intensity_StdIntensity_CMDR | 15 | 2678.51224 | | 0.0180 |
| GFP_Core_AreaShape_Compactness | 10 | 2157.85455 | | 0.0145 |
| spheroids_RadialDistribution_MeanFrac_GFP_3of6 | 11 | 2132.59715 | | 0.0143 |
| spheroids_Intensity_MeanIntensityEdge_GFP | 7 | 2081.51307 | | 0.0140 |
| Area_Ratio | 8 | 2047.6948 | | 0.0137 |
| GFP_Core_Intensity_StdIntensityEdge_CMDR | 12 | 1896.68842 | | 0.0127 |
| spheroids_Intensity_UpperQuartileIntensity_GFP | 5 | 1704.2935 | | 0.0114 |
| GFP_Core_AreaShape_FormFactor | 13 | 1668.01416 | | 0.0112 |
| GFP_Core_AreaShape_Solidity | 13 | 1513.71121 | | 0.0101 |

**E**

| Term | Splits | SS | | Portion |
|---|---|---|---|---|
| spheroids_RadialDistribution_MeanFrac_GFP_2of6 | 71 | 32492.9856 | | 0.1467 |
| spheroids_RadialDistribution_FracAtD_GFP_2of6 | 32 | 15031.9477 | | 0.0679 |
| spheroids_RadialDistribution_MeanFrac_GFP_1of6 | 39 | 12804.953 | | 0.0578 |
| GFP_Core_Intensity_IntegratedIntensityEdge_CMDR | 39 | 5470.25544 | | 0.0247 |
| spheroids_RadialDistribution_MeanFrac_GFP_4of6 | 42 | 4759.19859 | | 0.0215 |
| spheroids_RadialDistribution_FracAtD_GFP_4of6 | 32 | 2740.99846 | | 0.0124 |
| spheroids_Intensity_MADIntensity_GFP | 25 | 2356.75866 | | 0.0106 |
| spheroids_AreaShape_Solidity | 25 | 2230.59425 | | 0.0101 |
| spheroids_RadialDistribution_FracAtD_GFP_1of6 | 11 | 1644.79957 | | 0.0074 |
| spheroids_RadialDistribution_RadialCV_GFP_5of6 | 24 | 1545.12836 | | 0.0070 |
| spheroids_RadialDistribution_ZernikeMagnitude_GFP_6_0 | 23 | 1540.73961 | | 0.0070 |
| spheroids_RadialDistribution_ZernikeMagnitude_GFP_8_2 | 20 | 1487.50893 | | 0.0067 |
| GFP_Core_AreaShape_SpatialMoment_2_3 | 10 | 1468.71066 | | 0.0066 |
| GFP_Core_AreaShape_InertiaTensor_0_0 | 12 | 1415.2474 | | 0.0064 |
| spheroids_RadialDistribution_ZernikeMagnitude_GFP_2_0 | 15 | 1325.83756 | | 0.0060 |

**Supplemental Figure 4.** Radial distribution found as most instructive feature to determine SHROOM3 genotype percentage using either f-actin or ZO1 staining. (**a**) Specific details and results for the random forest predictive model utilized with the automated data presented in Figure 6. (**b**) Actual treatment concentration vs. predicted XY scatter-plot based on the bootstrap random forest model for the training data (left: 80% of dataset) and validation data (right: 20% of dataset) for the SHROOM3 SOSRS stained for f-actin (phalloidin-Alexa488). (**c**) Actual treatment concentration vs. predicted XY scatter-plot based on the bootstrap random forest model for the training data (left: 80% of dataset) and validation data (right: 20% of dataset) for the SHROOM3 SOSRS stained for ZO1. (**d**) Top 15 most instructive features (of 407 total) for the prediction of SHROOM3 genotype percentage using f-actin dataset. (**e**) Top 15 most instructive features (of 407 total) for the prediction of SHROOM3 genotype percentage using ZO1 dataset.
